# A proteomic perspective and involvement of cytokines in SARS-CoV-2 infection

**DOI:** 10.1101/2021.12.06.471525

**Authors:** Sarena Banu, Mohammed M Idris, Ramakrishnan Nagaraj

## Abstract

Infection with the SARS-CoV-2 virus results in manifestation of several clinical observations from asymptomatic to multi-organ failure. Biochemically, the serious effects are due to what is described as cytokine storm. The initial infection region for COVID-19 is the nasopharyngeal/oropharyngeal region which is the site where samples are taken to examine the presence of virus. We have earlier shown that several defensin genes are down regulated in cells from this region in patients who tested positive in the RTPCR test. We have now carried out detailed proteomic analysis of the nasopharyngeal/oropharyngeal swab samples collected from normal individuals and those tested positive for SARS-CoV-2 by RTPCR, involving high throughput quantitative proteomics analysis. Several proteins like annexins, cytokines and histones were found differentially regulated in the host human cells following SARS-CoV-2 infection. Genes for these proteins were also observed to be differentially regulated when their expression was analyzed. Majority of the cytokine proteins were found to be up regulated in the infected individuals. Cell to Cell signaling interaction, Immune cell trafficking and inflammatory response pathways were found associated with the differentially regulated proteins based on network pathway analysis.

## Introduction

The COVID19 pandemic has lead to extensive investigations on multiple aspects of the biology of SARS-CoV-2 virus as well as host-responses [1,2]. Despite extensive investigations, as of now, vaccination appears to be the only way to avoid or reduce infection by the virus [1, 2]. To date, testing for the virus relies on swabs from nasopharyngeal region followed by detection of the virus by RTPCR [1-3]. There have been detailed proteomics studies on the serum of patients infected with SARS-CoV-2 [4-8]. These studies indicate up or down regulation of several proteins on infection in the serum. It is important to investigate early events at the initial site of infection immediately after detection by RTPCR. The early events at the site of infection, may give a clue about the disease progression and the protein regulation at an early stage of infection.

The initial site of infection by the virus is the nasopharyngeal region [1-4, 9-11]. Hence, it is relevant to examine the effect of virus attack on the cells of this region. We have shown that upon infection, there is down regulation of defensin genes [12]. Defensins are crucial components of innate immunity and are the first line of defense against invading pathogens [12, 13]. In the present study, we have now analyzed the cells from this region for expression of protein profiles, particularly their regulation with respect to controls. We have also in parallel examined differential expression of different cytokines. We have observed up regulation of several proteins related to inflammation and innate immunity. Of particular interest is the up regulation of annexins. Auto antibodies to annexins have been observed in covid19 patients [14]. The up regulated proteins may result in the production of auto antibodies which contributes to metabolic dysfunction observed in the disease. We have observed up regulation of several cytokines at the genomic and protein level. These cytokines play a role in inflammation. Our results indicate that metabolic stress due to viral infection is observed in nasopharyngeal cells at the onset of infection. Hence, early therapeutic intervention could be beneficial as catastrophic metabolic dysfunction such as cytokine storm could be avoided.

## Results

The Nasopharyngeal region comprises of a heterogeneous population of cells that include epithelial and immune cells [15, 16]. The Nasopharyngeal region is also gateway to the lungs which is affected badly when the illness is serious. The detection of SARS-CoV-2 infection involves RTPCR from Nasopharyngeal/oropharyngeal swab (NOPS) samples. We have earlier shown that analysis of cells from this region indicates down regulation of defensin gene expression [12]. We describe in this paper, differential proteomics analysis and expression of some cytokine genes from NP swabs from control population and those infected with SARS-CoV-2 as indicated by RTPCR.

### Proteomics analysis

Six major proteins important for virus entry into cells replication such as ORF1ab, N, nsp9, S, nsp3 and H [1] were found expressed in the infected NOPS following high throughput proteomic analysis (Table 1). A total of 37 proteins of host humans were found differentially regulated on infection with SARS-CoV-2 (Table 2). The differentially regulated proteins include 21 up regulated and one down regulated protein for having more than one log fold differential regulation (Table 2). Proteins such as HIST1H3A, H2AFZ, HIST1H4A, S100A9, GSTA1, DPYSL5 and S100A8 were found to be up regulated by 4 log fold changes. ANXA2, BPIFA1, AXNA1, HIST1H2BK and AKR1C4 were up regulated by 3 log fold change. Proteins that were up regulated 1-3 fold were: LTF, ATP5F1A, YWHAQ, ANXA5, LYZ, PRDX1, DMBT1, LGAL53 and CAPS (Table 2). Many of the up-regulated proteins are involved in host-defense functions and inflammatory reactions.

**Table 1:**
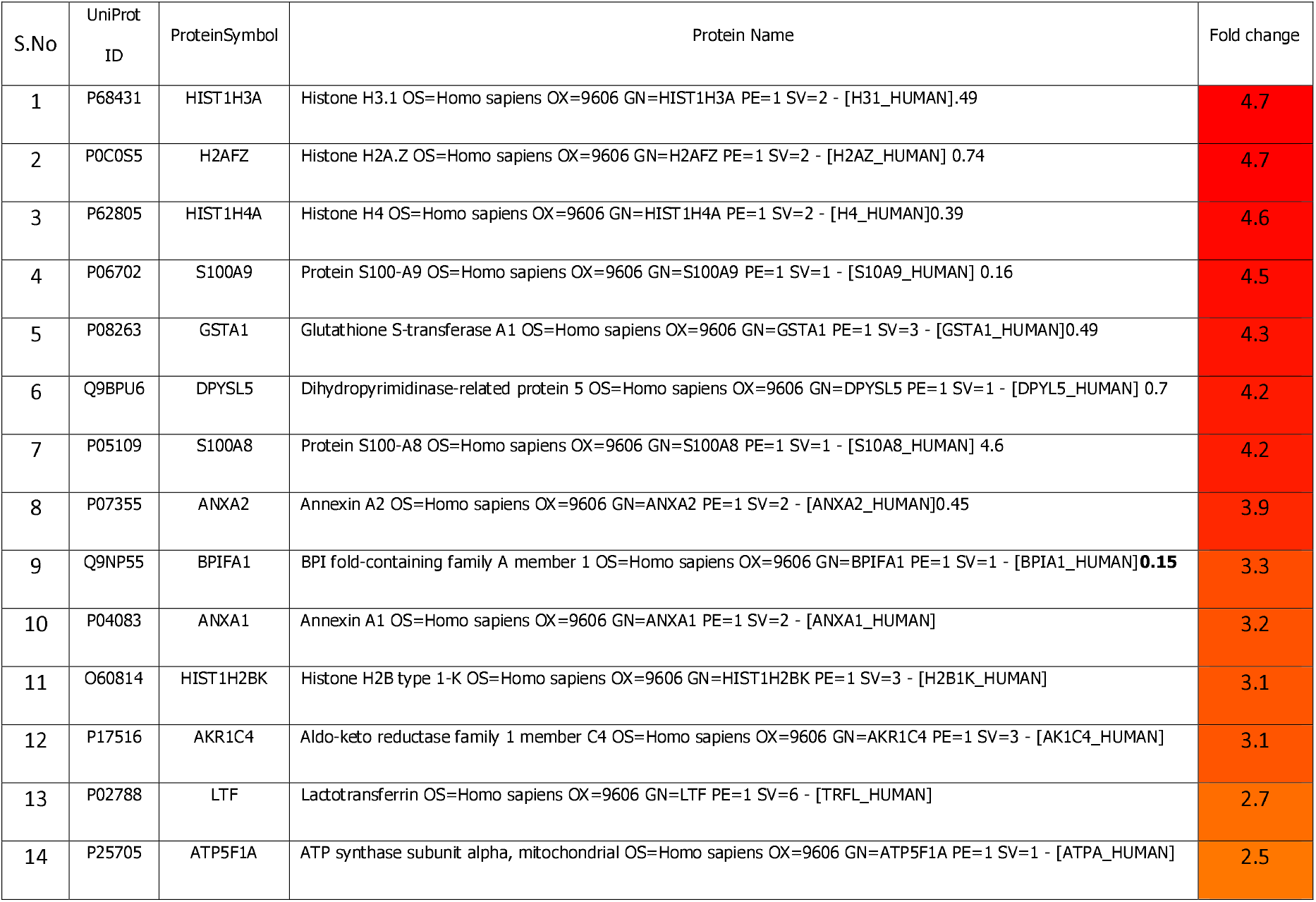
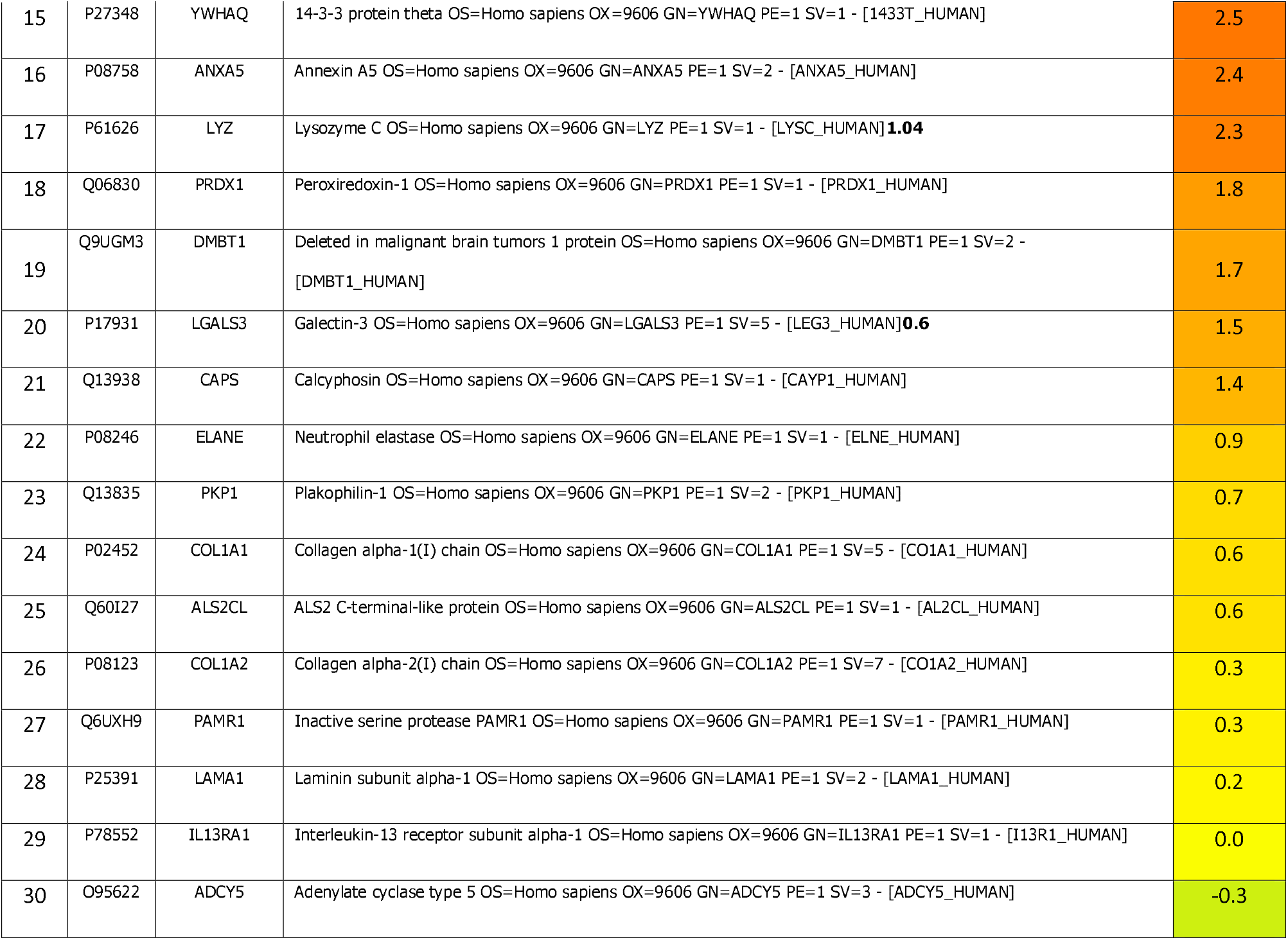
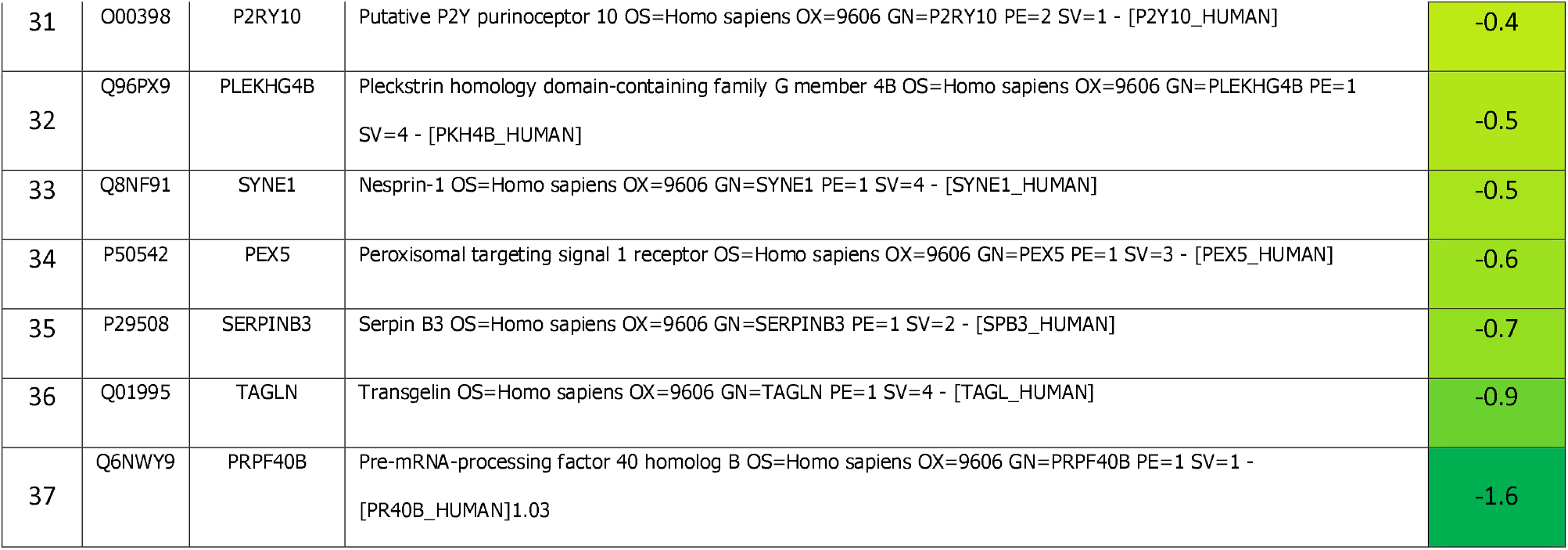
List of proteins which were found differentially regulated in the infected NOPS samples against control negative samples

**Table 2:** List of viral proteins which were found expressed in the COVID infected NOPS samples.

| S.No | UniProt ID | Protein<br>Symbol | Protein Name |
| --- | --- | --- | --- |
| 1 | YP_009724389.1 | ORF1ab | ORF1ab polyprotein [Severe acute respiratory syndrome coronavirus 2] |
| 2 | YP_009724397.2 | N | nucleocapsid phosphoprotein [Severe acute respiratory syndrome coronavirus 2] |
| 3 | YP_009742616.1 | nsp9 | nsp9 [Severe acute respiratory syndrome coronavirus 2] |
| 4 | YP_009724390.1 | S | surface glycoprotein [Severe acute respiratory syndrome coronavirus 2] |
| 5 | YP_009742610.1 | nsp3 | nsp3 [Severe acute respiratory syndrome coronavirus 2] |
| 6 | YP_009725308.1 | H | helicase [Severe acute respiratory syndrome coronavirus 2] |

### Gene expression analysis

Genes of proteins involved in immune responses [17] and those that were observed to be up regulated in the proteomics analysis were analyzed for their expression involving RTPCR during SARS-CoV-2 infection using respective gene specific primers (Table 3). Based on RTPCR analysis it was found that majority of the selected genes were significantly up regulated on viral infection. IL1b, IL11, IL6, ICAM1, VCAM1, HIST1H2BK, HIST1H4A, H2AFZ, HIST1H3A, SERPINB3, DMBT1, ANXA1, ANXA2 were found up-regulated by one log fold change (Figure 1). LYZ is the only gene which is found down regulated upon SARS-CoV-2 infection (Figure 1). Proteins which were found up regulated were also up regulated at the gene level through RTPCR.

**Table 3:**
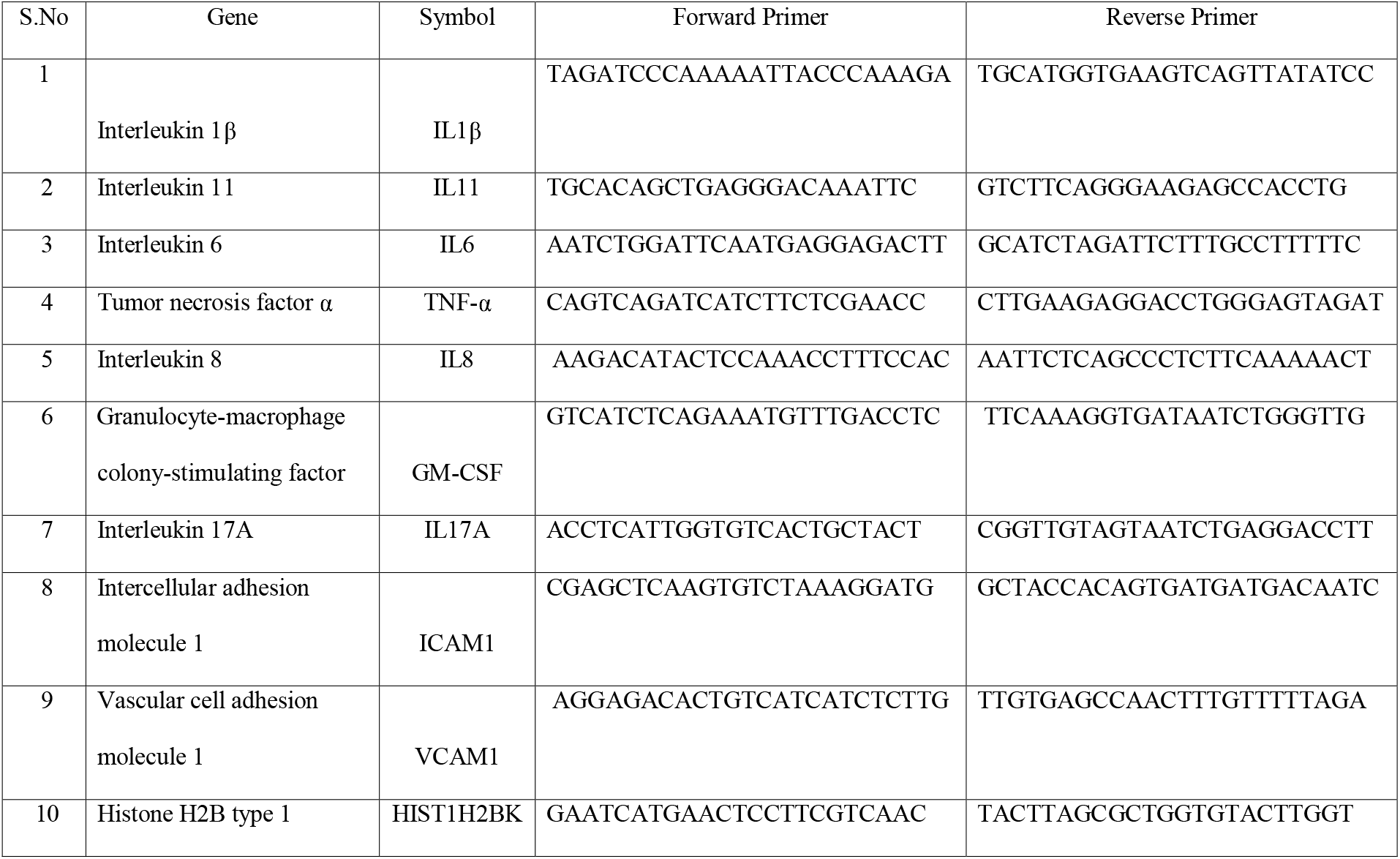
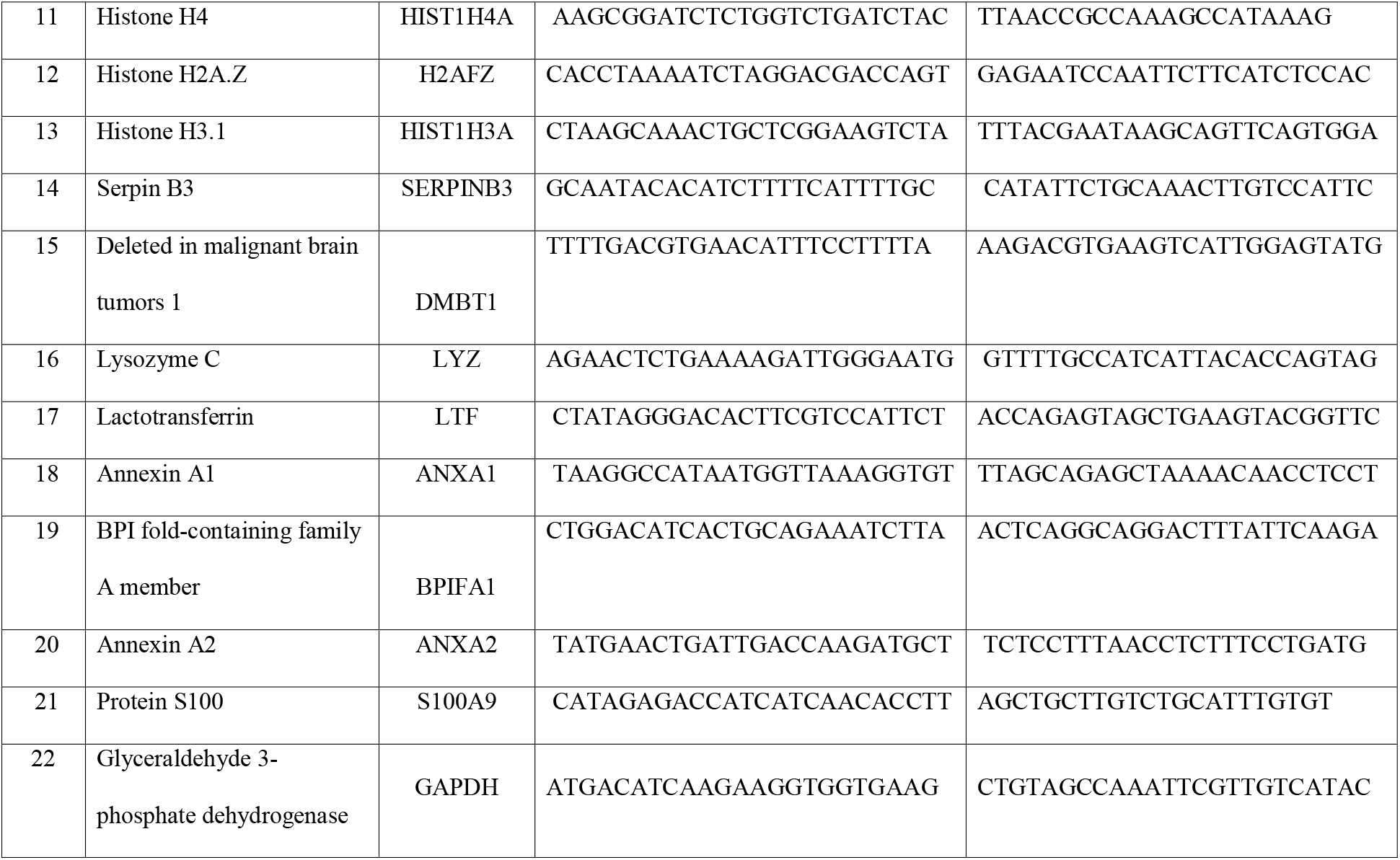
List of genes and their primer pairs used for the RTPCR validation

**Figure 1:**
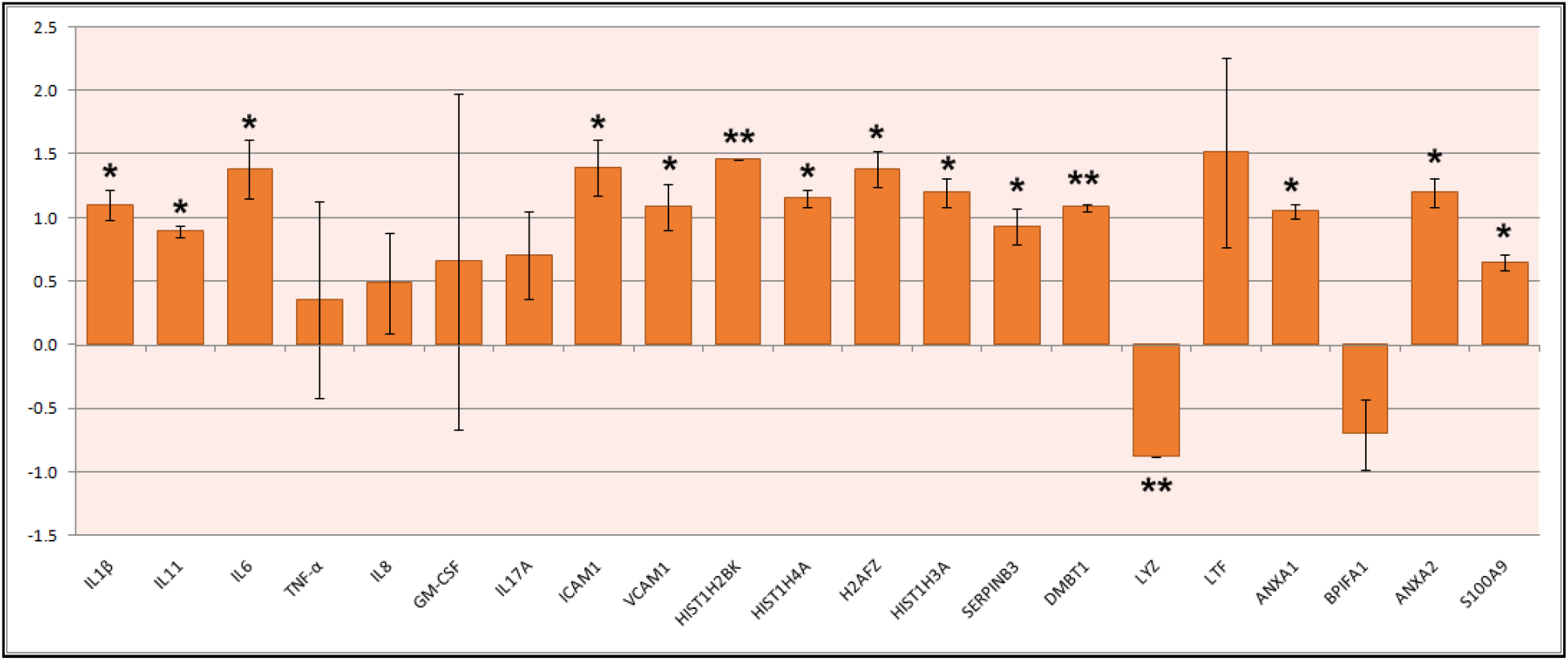
qRTPCR expression analysis of genes associated with infection. The bar represents the relative expression of the gene in SARS-CoV-2 infected samples against negative controls.

### Network Pathway Analysis

Based on network pathway analysis it was found that inflammatory response network pathway was highly associated with the differentially regulated proteins and genes observed and selected in the study. Genes/proteins (16 in number) which were found differentially regulated in the NOPS samples for SARS-CoV-2 infection were found associated with inflammatory response pathway (Figure 2a). Cellular movement/cell to cell signalling and cellular assembly/organization were the two major canonical pathways found to be associated with the differentially regulated proteome of infected NOPS (Figure 2b). The major functions associated with the identified and differentially regulated proteome includes cell to cell signalling and interaction, immune cell trafficking, cellular movement, inflammatory response and hypersensitive response (Figure 3).

**Figure 2:**
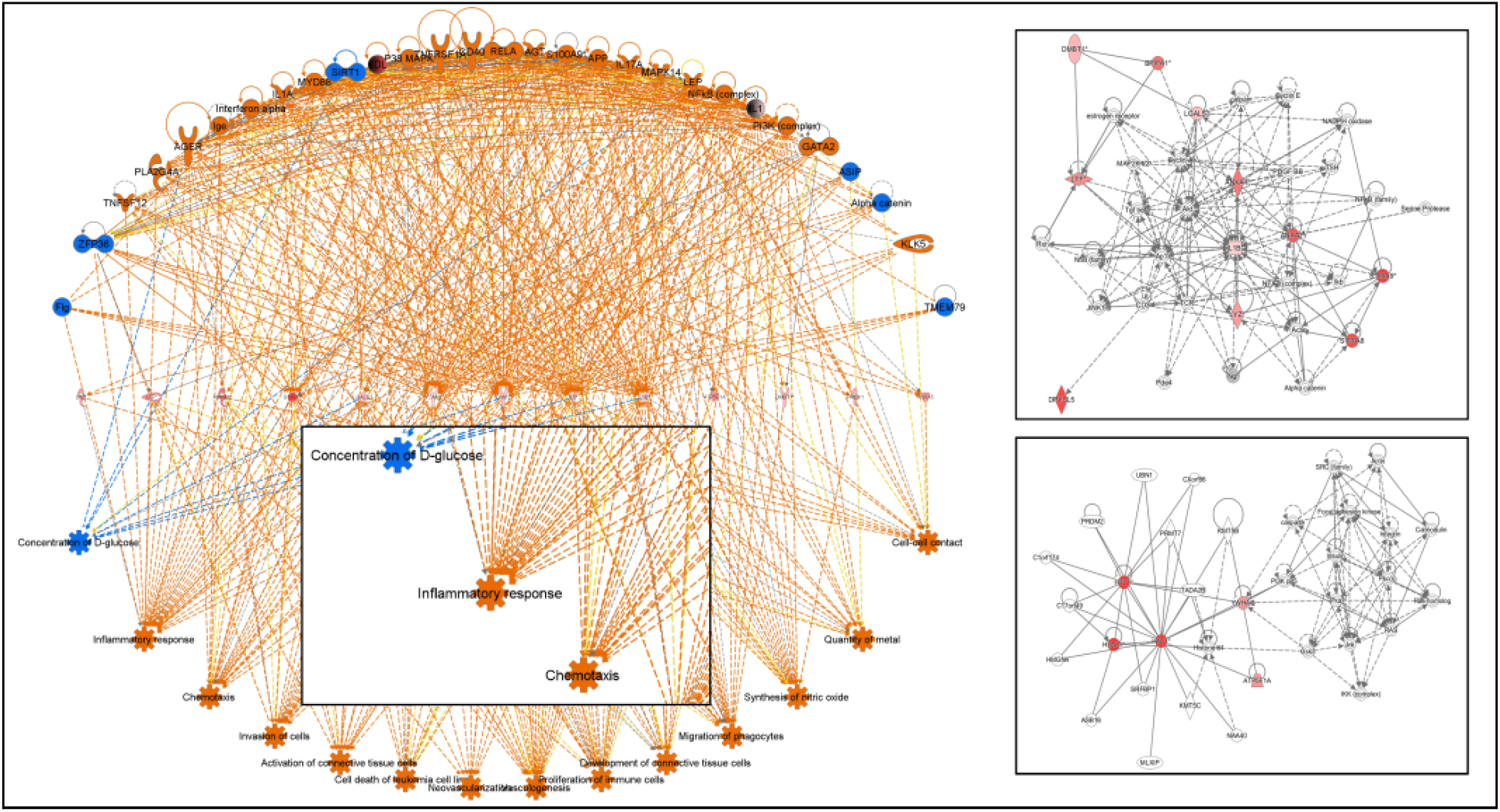
Network pathway analysis of the genes/proteins associated with infection in NOPS samples. a. Interaction network based on IPA analysis. b. Two most significant network associated with infection.

**Figure 3:**
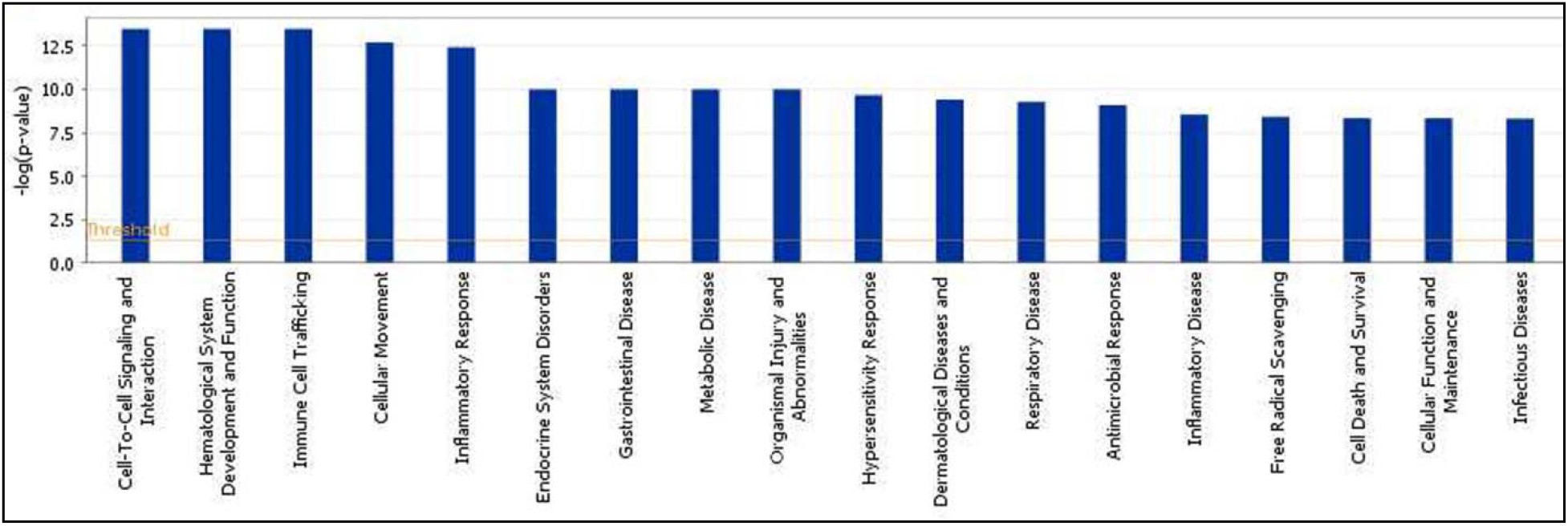
List of functions (or disease) associated with the differentially expression proteins/genes which are associated with infection.

### Cytokine array and western blot analysis

Based on cytokine array analysis it was found that almost all the proteins represented for the cytokine pathway [17] were found to be up regulated in the infected Nasopharyngeal region samples against the control NOPS samples. Majority of the interleukins, VEGF, CNTF and CINC-2 were found to be majorly up regulated for COVID infection based on cytokine array analysis. Based on western blot analysis it was found that ANXA1, ANXA2 and ANXA5 were found to be up regulated by several fold in SARS-CoV-2 positive NOPS samples against control NOPS samples (Figure 5). ANXA2 was found up regulated majorly in the infected samples.

## Discussion

The initial site of SARS-COV-2 infection is the upper airways. As the disease progresses, respiratory difficulties have been observed in severe cases [2]. The expression of two main SARS-COV-2 receptors, ACE2 and TMPRSS, was found to be high in the upper airway tract mucosa [15]. The nasopharyngeal region plays a major role in mucosal immunity against SARS-COV-2 infection by activating lymphocytes, B cells, and other cellular components, as well as activating antigen specific immunity [9]. The oral cavity was also reported to be implicated in various immunological responses to SARS-COV-2 infection [15]. In fact, the test for viral infection is from swabs from this region which are subjected to RTPCR. The swabs would contain epithelial and immune cells [16, 18]. We have shown that when cells from this region were analyzed for the expression of host-defense peptides-the defensins, they were significantly down-regulated [12]. In this paper, we described the effect of SARS-CoV-2 infection in patients on the expression of cytokine genes and proteomics profile of cells from the nasopharyngeal region.

Global proteome studies were done on infected SARS-COV-2 *in vitro* cells [19], serum samples [4, 7], blood samples from COVID-19 patients [8] and nasopharynx mucosa [20] to study the host response to SARS-COV-2 infection. In this study we have analyzed the proteomic changes in the nasopharyngeal region following SARS-CoV-2 infection involving high throughput quantitative differential proteomic analysis.

We have analyzed samples from individuals who were tested COVID19 positive by RTPCR test and individuals who were COVID19 negative. The major viral proteins were detected in the infected nasopharyngeal samples. They were SARS-CoV-2non-structural proteins that are responsible for viral transcription, replication, proteolytic processing, suppression of host immune responses and suppression of host gene expression (Table 1). The nucleocapsid protein is an RNA-binding protein that is essential for viral assembly into a ribonucleoprotein complex and also functions in viral budding.

The host proteins which are found to be differentially regulated for the SARS-CoV-2 infections are found to be involved in various aspects of cell signalling and immune response. HIST1H3A, H2AFZ, HIST1H4A, GSTA1, DPYSL5, S100A9, and S100A8 proteins were found to be up regulated by more than 4 log fold changes. It is interesting that there is up-regulation of histones, which are DNA-binding proteins, on infection. Histones have a significant role in inflammation (Table 2). When compared to other histones, H3 and H4 have the strongest antiviral action against influenza A viruses. Histones are also a prominent component of neutrophil extracellular traps (NETs), which causes inflammation. Histones at high concentrations (50ug/ml) have been linked to lung damage [21]. Up regulation of histone proteins may play a role in host innate immune modulation as well as severe inflammatory reactions such as host epithelial and lung endothelial cell injury.

SARS-COV-2 infection causes oxidative stress in host cells and stimulates the production of cytokines, antioxidants, transcription factors, neutrophils, dendritic cells, and macrophages [22]. Glutathione S transferase is a detoxifying enzyme that protects host cells from oxidative stress caused by SARS-COV-2 [22]. GSTA1 down regulation in human lung adenocarcinoma cell line A549 promotes apoptosis, which suppresses tumor development [23].Glutathione S-transferase host oxidative defensegeneGSTT1 and GSTM1 polymorphism was observed in SARS-COV-2 infection [22].Peroxiredoxin-1 (PRDX1), an antioxidant, also protects against oxidative stress [24]. Lysozyme C is an antibacterial enzyme that inhibits oxidative stress and inflammation as a result of viral infection [4, 25]. It also acts as an immunological modulator. Our findings show that GSTA1, PRDX1, and LYZ proteins have a significant oxidative defense mechanism during SARS-COV-2 infection.

S100A8/A9 or calprotectin is a neutrophil activated protein; high levels of S100A8/A9 expression were found in COVID-19 severe patients’ blood samples [26]. It is noteworthy to explore anti-neutrophil therapy in context of significant up regulation of the S100A8/A9 gene and protein expression. Upregulation of the DPYSL5 protein suggests that SARS-CoV-2 infection causes immune-mediated neuropathies [27]. Our proteomics study showed that the protein neutrophil elastase (ELANE) was up regulated. Neutrophil elastase (ELANE) promotes acute lung injury by activating pro-inflammatory cytokines such as IL-6 and IL-8 [28]. It has been found that inhibiting neutrophil elastase levels reduces acute lung injury [28]. ELANE may also be one of the critical factors causing a severe lung infection in SARS-CoV-2 infections.

Our proteomic findings are also consistent with previous *in vitro* studies that showed SARS-CoV-2 infection activated host immune proteins. The SERPINB3 gene was found to have high expression in the upper airway tract [29]. SERPINB3 stimulates NF-B signalling and the pro-inflammatory response, specifically IL-6 [29, 30]. IL-6can bind to TMPRSS2 and prevent SARS-CoV-2 entrance into the host cell [29].Deleted in malignant brain tumors 1 (DMBT1) binds to several growth factors such as VEGF and FGF and inhibits IL-6 production in lung epithelial cells [31]. Lactotransferrin (LTF) similarly inhibits IL-6 production which has been shown to be antiviral in SARS-COV-2 infections [32].

Several other proteins as described in Table 2 are also associated with inflammatory responses and apoptosis. Of particular interest is the up regulation of Annexins (Figures 1 and 5) (Table 2). Annexin family proteins are calcium dependent phospholipid binding proteins. Annexin A1 (ANXA1), Annexin A2 (ANXA2), Annexin A5 (ANXA5) predominantly found in SARS-CoV-2 infected host cells [33]. ANXA1, 2, and 5 are involved in the pro-inflammatory response and thrombosis [33, 34]. Recent reports indicate that auto-antibodies to Annexin 2 is detected in patients with COVID19 [14, 34].

Up regulation of cytokines based on cytokine protein array (Figure 4) analysis directly correlates the association of inflammatory responses to infection. Both pro-and anti-inflammatory cytokines and several growth factors involved in inflammatory response were found up regulated for the infection. Cytokines are a major component of the innate immune response and are produced by lymphocytes and granulocytes [35-37]. During inflammatory reaction, Host cells pathogen-recognition receptors (PRRs) triggers cytokines in host system [19, 35-37]. It is divided into two types: pro-inflammatory cytokines, which release excessive cytokines in response to infection, and anti-inflammatory cytokines, which regulate pro-inflammatory cytokines [35-37]. Pro-inflammatory cytokines such as IL-1, IL-10, IL-6, and TNF-α were found in significant concentrations in COVID-19 individuals [36, 38]. Pro-inflammatory cytokines cause a cytokine storm during SARS-CoV-2 infection, which has been linked to ARDS, multi-organ failure, and increased candidate risk [36, 38]. Our study found that pro-inflammatory cytokines such as IL-1, IL-6, IL-10, and TNF-α, as well as other cytokines, were highly expressed in COVID-19 patients at the protein level (Figure 4). At genomic expression, IL-6 found to be significantly up regulated with more than 1 log fold change (Figure 1).Other cytokines such as IL-2,IL4, GM-CSF, IFN-γ,MCP-1 also found to be associate with SARS-COV-2 infection [36].The pro-inflammatory cytokine TNF-α, regulates leukocyte trafficking by stimulating cell adhesion molecules such ICAM-1, VCAM-1, and selectins [39]. VCAM-1 plays an important role in leukocyte trafficking through vascular adhesion and other mechanisms. VCAM-1 has also been associated with asthma, rheumatoid arthritis, and a number of immune-related diseases [39]. Our data showed that ICAM-1 and VCAM-1 gene (Figure 1) up regulated with more than one log fold change when compared to COVID-19 negative patients. It implies that VCAM-1 could be exploited as a therapeutic target. ICAM-1 and L-selectin protein also showed up regulation with more than 1 log fold in cytokine array (Figure 4). The expression level of cytokines was examined using the probes is detailed in Table 3. Enhanced expression of cytokines is observed. The network path analysis indicates links between cell-cell signalling and immune response (Figure 2).

**Figure 4:**
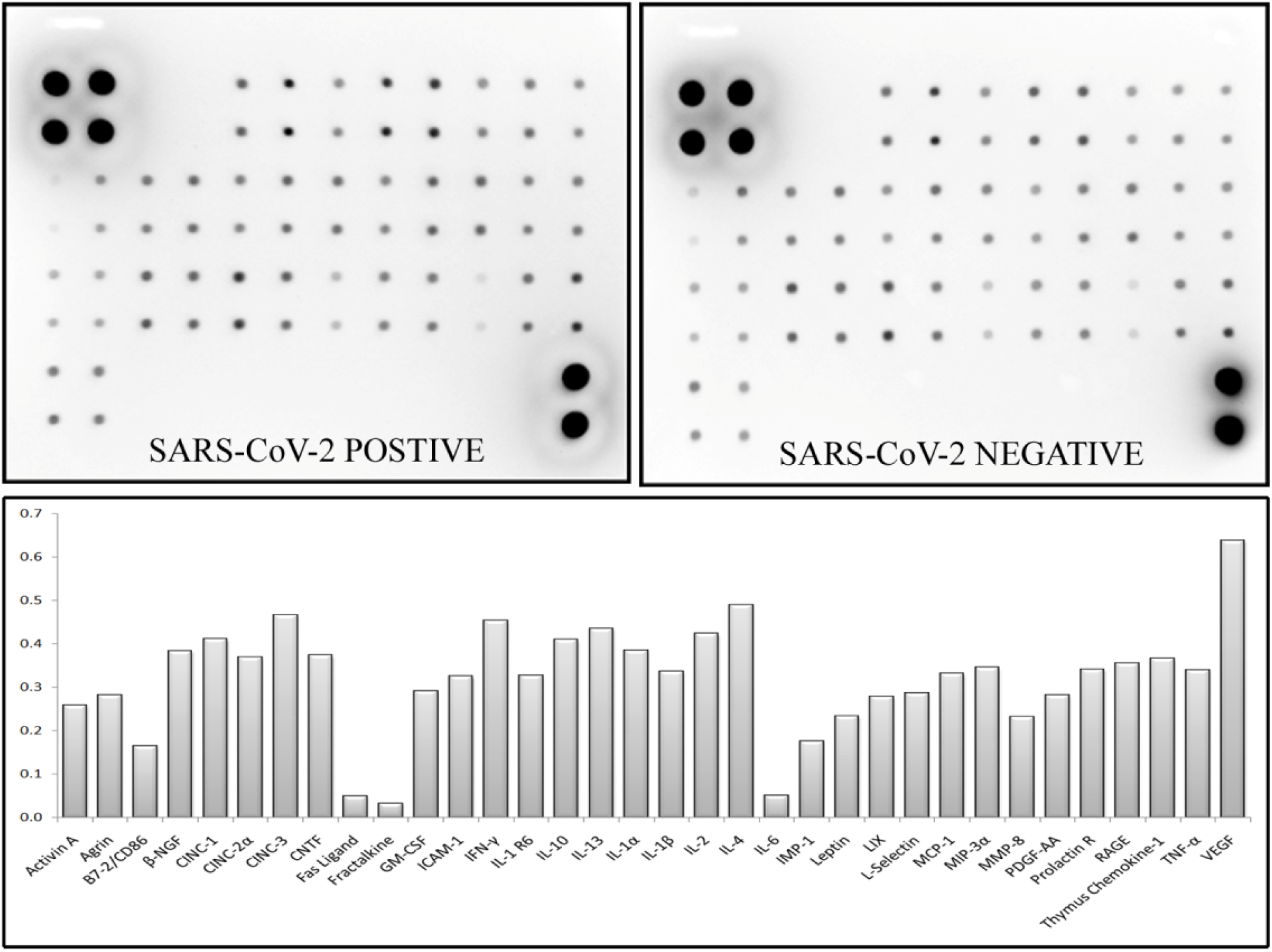
Cytokine protein array analysis for SARS-CoV-2 positive and negative samples. Bottom panel represent the relative expression of the protein based on cytokine protein array analysis.

**Figure 5:**
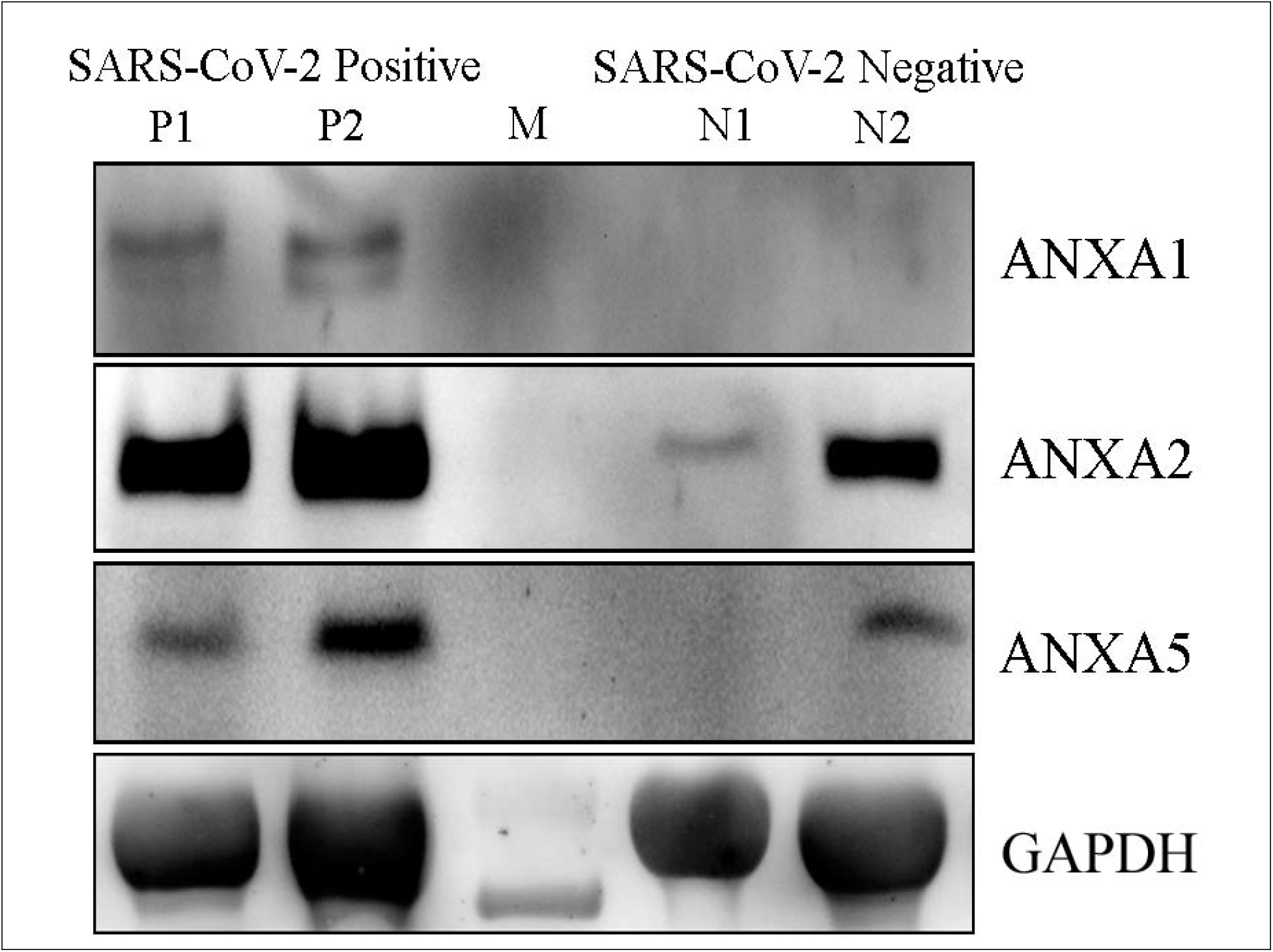
Western blot analysis for ANXA1, ANXA2 and ANXA5 protein in infected and non-infected NOPS samples. P1 and P2 indicates the SARS-CoV-2 positive samples; M denotes the protein molecular weight standard and N1 and N2 denotes the SARS-CoV-2 negative samples..

Although infection by corona virus was manifested in a mild form earlier on [1,2], the pandemic due to the corona virus SARS-CoV-2 that started in 2019 had a devastating effect on health services globally. Despite extensive efforts, there is no drug specifically to treat COVID19. There have been extensive researches on various aspects on the virology of SARS-CoV-2, not only to understand the disease, but also to devise effective treatment. Development of vaccines at warp speed has been effective in controlling the disease [40], but data on long term protection are not available. The symptoms are highly variable among infected individuals and no generalization is possible. Responses by various populations may differ considerably. Hence, it is important to investigate protein and gene expression from patients from different geographical locations and ethnicity. It is important to examine various biochemical parameters on the onset of infection particularly from the initial site of infection to understand the disease. By a combination of proteomics analysis, gene expression studies as well as in the blot array, we were able to detect several proteins related to inflammation and auto immunity such as histones and annexins and up regulated cytokines. Clearly a multi-dimensional approach is necessary to obtain a complete picture on SARS-CoV-2 infection. Such an approach would facilitate therapeutic interventions.

## Methods

### 1. Sample collection

Human SARS-CoV-2 viral transport media (VTM) samples were obtained from CSIR-CCMB COVID19 diagnostic facility [12]. A total of 80 VTM samples (collected from Government hospital, Hyderabad during early 2020) grouped into 8 with each group consisting of 5 positive and 5 negative VTM samples were taken for the gene expression study. A total of 12 VTM samples grouped into 2 with each group consisting of 3 positive and 3 negative SARS-CoV-2 VTM samples were taken for the quantitative differential proteomics study. The study was executed as per the approval of CCMB Institutional biosafety and institutional ethical approval.

### 2. Protein Extraction and iTRAQ labelling

Total protein was extracted from the two groups consisting of 3 positive and 3 negative pooled SARS-CoV-2 VTM samples. At the BSL3 lab facility, the pooled samples were centrifuged at high RPM, and the resulting pellet was washed with 1% PBS, resuspended in protein solubilization buffer, and sonicated for 10 minutes. After brief centrifugation, the supernatant was collected and quantified using the amido black method [41]. A total of 200ug of protein was in-gel trypsin digested and labelled with iTRAQ label as per previously mentioned protocol [41-45]. SARS-COV-2 negative samples were labelled with iTRAQ 114 and 115, and positive samples were labelled with 116 and 117. All the labelled peptides were pooled and analysed using Liquid Chromatography Mass Spectrometry (LCMS/MSMS) in the Orbitrap Velos Nano analyzer (Q-Exactive HF). The obtained data were analyzed against the human proteome and SARS-CoV-2 proteome database. The identified SARS-COV-2 and host proteins were listed. Differential expression analysis was carried for the obtained host proteins against control SARS-CoV-2 negative samples.

### 3. Real-time PCR analysis

Total RNA was extracted from the selected individual VTM swab samples using KingFisherTM Flex System (Thermo Fisher Scientific Inc., USA).A total of 80 VTM RNA samples were quantified and grouped into 8 with each group consisting 100ng of 5 positive and 5 negative samples. cDNA was synthesized (BioRad, USA) from 200 ng of each VTM group. Real-time PCR analysis was performed for the selected list of the gene using Applied BiosystemsViiA™ 7 Real-Time PCR System (USA) with gene-specific primers (Table 3). All primers were synthesized using Primer3 software and, the GAPDH gene was used for the data normalization. TB Green Premix Ex Taq 11 (TliRNaseH Plus) kit (Takara, Japan) with respective qRTPCR conditions (Annealing Tc-55°C or 60°C) and melt curve analysis were followed as previously mentioned protocol [12]. Differential expression analysis was performed using the obtained cycle threshold value.

### 4. Cytokine array and western blot analysis

Cytokine expression analysis was performed using Cytokine Array - RAT Cytokine antibody array (Abcam, USA) kit. A total of 300 µg proteins from two groups consisting of 6 positive and 6 negative samples were immunoblotted in the kit as per the manufacturer’s protocol. The obtained spot patterns were densitometrically analyzed using ImageJ software to estimate the expression level of cytokines in SARS-CoV-2 infection.

Protein expression analysis of ANXA1, ANXA2 and ANXA5 protein were analyzed involving western blot analysis. 25 µg of total protein electrophoresed in 10% SDS-PAGE and, immune blotted against respective antibodies [46]. GAPDH was used as a housekeeping protein.

### 5. Pathway Analysis

The proteins and genes selected for the study were analyzed for network and pathway analysis using Ingenuity Pathway Analysis software-based. The network pathway, regulator effect and disease and function associated with the genes/protein were mapped.

## Author’s contribution

SB – Performed the experiments and analyzed the data; MMI – conceived, analyzed and wrote the MS; RN-conceived, analyzed and wrote the MS.

## Acknowledgements

RN is Indian National Science Academy Senior Scientist.

